# Consensus design and engineering of an efficient and high-yield Peptide Asparaginyl Ligase

**DOI:** 10.1101/2022.11.02.514816

**Authors:** Xinya Hemu, Xiaohong Zhang, Hong Yi Chang, Poh Jin En, James P. Tam

## Abstract

Plant legumains are Asn/Asp-specific endopeptidases (AEPs) that have diverse functions in plants. Peptide asparaginyl ligases (PALs) are a special legumain subtype that primarily catalyze peptide bond formation rather than hydrolysis. PALs are versatile protein engineering tools but are rarely found in nature. To overcome this limitation, here we describe a two-step method to design and engineer a high-yield and efficient recombinant PAL based on commonly found AEPs. We first constructed a consensus sequence derived from 1,500 plant legumains to design the evolutionarily stable legumain conLEG that could be produced in *E. coli* with 20-fold higher yield relative to that for natural legumains. We then applied the LAD (ligase-activity determinant) hypothesis to exploit conserved residues in PAL substrate-binding pockets and convert conLEG into conPAL1-3. Functional studies showed that conLEG is primarily a hydrolase, whereas conPALs are ligases. Importantly, conPAL3 is a super-efficient and broadly active PAL for peptide and protein cyclization.

## Introduction

Legumains are cysteine proteases that are widely distributed among plants and animals. These proteases were named after their discovery in legume seeds in the early 1990s (Kembhavi et al., 1993). They have also been isolated from parasites (Dalton et al., 1995) and mammals (Chen et al., 1997). Legumains belong to the same family of enzymes as vacuolar processing enzymes (VPEs) that were discovered in the late 1980s (Hara-Nishimura & Nishimura, 1987). Regardless of their origin, these enzymes share a similar protein fold and are classified in the C13 subfamily of Cys proteases (MEROPS, EC 3.4.22.34) (Rawlings et al., 2018). Functionally, legumains are asparaginyl endopeptidases (AEPs) that hydrolyze the peptide bond after an Asn/Asp(Asx) (Figure 1). Mammalian legumains are lysosomal enzymes that play important roles in antigen-presentation and cancer-related events (Dall & Brandstetter, 2016). In contrast to animal genomes that have only one gene encoding a legumain, plant genomes contain multiple copies of legumains. The major functions of plant legumains are to regulate senescence processes (Hatsugai et al., 2015), maturation of seed storage proteins (Gruis et al., 2004), and processing of defensive proteins and peptides.

**Figure 1.**
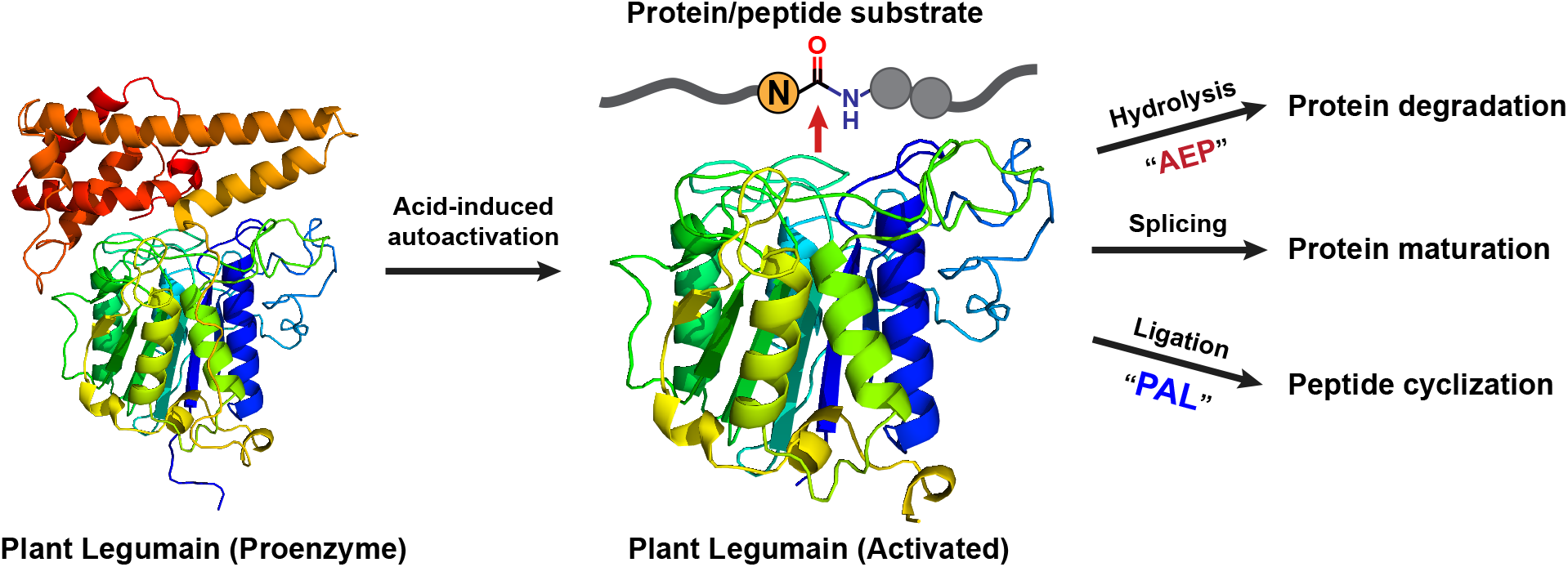
Schematic illustration of activation, enzymatic activity and biological functions of plant legumain.

In addition to cutting peptides at specific Asx sites, legumains can also join peptides. An early example of this activity was observed in the maturation of the legume lectin, concanavalin A (conA), which involves a splicing (cut-and-join) mechanism. This protease-mediated peptide ligation is assisted by the correctly folded conformation of the conA precursor that brings two ligating termini in close proximity to facilitate a transpeptidation reaction. Recently, we identified a subtype of plant legumains that act as ligases rather than proteases. We termed these enzymes peptide asparaginyl ligases (PALs). Examples of PALs include the prototype, butelase-1 from *Clitoria ternatea* (Nguyen et al., 2014), which has a catalytic efficiency of 1.3×10^6^ M^-1^s^-1^, making it the fastest natural peptide ligase (Nguyen et al., 2015). Other PALs include OaAEP1b from *Oldenlandia affinis* (Harris et al., 2015; Harris et al., 2019), and a dozen more from *Viola* plants (Hemu et al., 2019; Jackson et al., 2018; Rajendran et al., 2021). Compared with widely distributed AEPs in both primitive and higher plants, PALs in nature are much rarer, only accounting for 1% of sequenced plant legumains (Hemu et al., 2022).

Butelase-1 and other PALs serve as bioprocessing enzymes of macrocyclic peptides such as cyclotides and trypsin inhibitors (Bernath-Levin et al., 2015; Du et al., 2020; Harris et al., 2015; Hemu et al., 2019; Liew et al., 2021; Nguyen et al., 2014). They recognize an Asx-Xaa-Yaa tripeptide motif in which Xaa represents any residue except Pro, and Yaa represents hydrophobic residues (Hemu et al., 2019; Nguyen et al., 2014). PALs function as efficient, site-specific ligases that do not require ATP for activity. As such, they are invaluable for biochemical and pharmaceutical applications for protein engineering and site-specific modifications of proteins and live cells (Bi et al., 2017; Cao et al., 2015; Harmand et al., 2018; Hemu et al., 2016; Hemu et al., 2021; Nguyen et al., 2016; Rehm et al., 2019).

A challenge to the wider use of PALs in industrial applications is associated with their recombinant productions. Recombinant PAL proenzymes can be expressed in bacteria, but the expressed proteins are often present as misfolded proteins in inclusion bodies (Nguyen et al., 2014) and the yield of soluble proteins is fairly low (Tang & Luk, 2021). Although incorporation of large fusion tags such as MBP can increase the yield of crude fusion proteins, they can also lead to heterogeneous activity of the enzymes after activation (Zhao et al., 2021; Zhao et al., 2022). Secretory expression in insect cells could produce about 20 mg/L of VyPAL2, but this approach is limited by relatively high costs. Similarly, secretory expression of C-terminal truncated butelase-1 in yeast yielded 16 mg/L active enzyme (Pi et al., 2019), but the success of this method has yet to be repeated for production of other PALs (data not shown). Attempts to improve expression levels using disulfide-promoting strains, codon modification, truncation of the flexible N-terminal pro-region, and incorporation of fusion tags suggest that expression of soluble PALs could be facilitated by increasing the stability of either cDNA, mRNA, or peptide chains. By using residues that appear most frequently among homologous genes, an engineered functional protein could acquire enhanced stability, or activity, or both (Gomez-Fernandez et al., 2020; Lehmann et al., 2002; Lehmann et al., 2000; Qian et al., 2018; Yao et al., 2020).

Here, we report a strategy to design high-yield PALs. We developed this strategy using bioinformatic analysis of a dataset containing 1,500 plant legumains to build a consensus legumain followed by mutation of substrate-binding pocket residues based on the LAD (ligase-activity determinants) hypothesis that was previously developed to convert an AEP to a PAL.

## Results

### Consensus plant legumain (conLEG) sequence for recombinant protein expression

As recently reported, we prepared a dataset containing 1,500 plant legumains from 249 plant families was constructed based on BLAST searches of existing protein sequences and transcriptomes in NCBI and 10KP databases, respectively, using representative AEPs and PALs as queries (Hemu et al., 2022). Multiple sequence alignment of these legumains produced a consensus sequence having a typical protein fold of C13 Cys proteases, including an α56 core-domain carrying the “Asn-His-Cys” catalytic triad and a 5-helix cap-domain (Dall & Brandstetter, 2013) (Figure S1).

The expression construct conLEG was designed by removing residues with <10% occupancy, substituting the signal peptide and non-conserved region of the pro-domain with a His6 affinity tag, and substituting the C-terminal unstructured loop after helix α10-Pro457 (corresponding to butelase-1 Pro469) with a C-terminal His6-tag (Figure 2).

**Figure 2.**
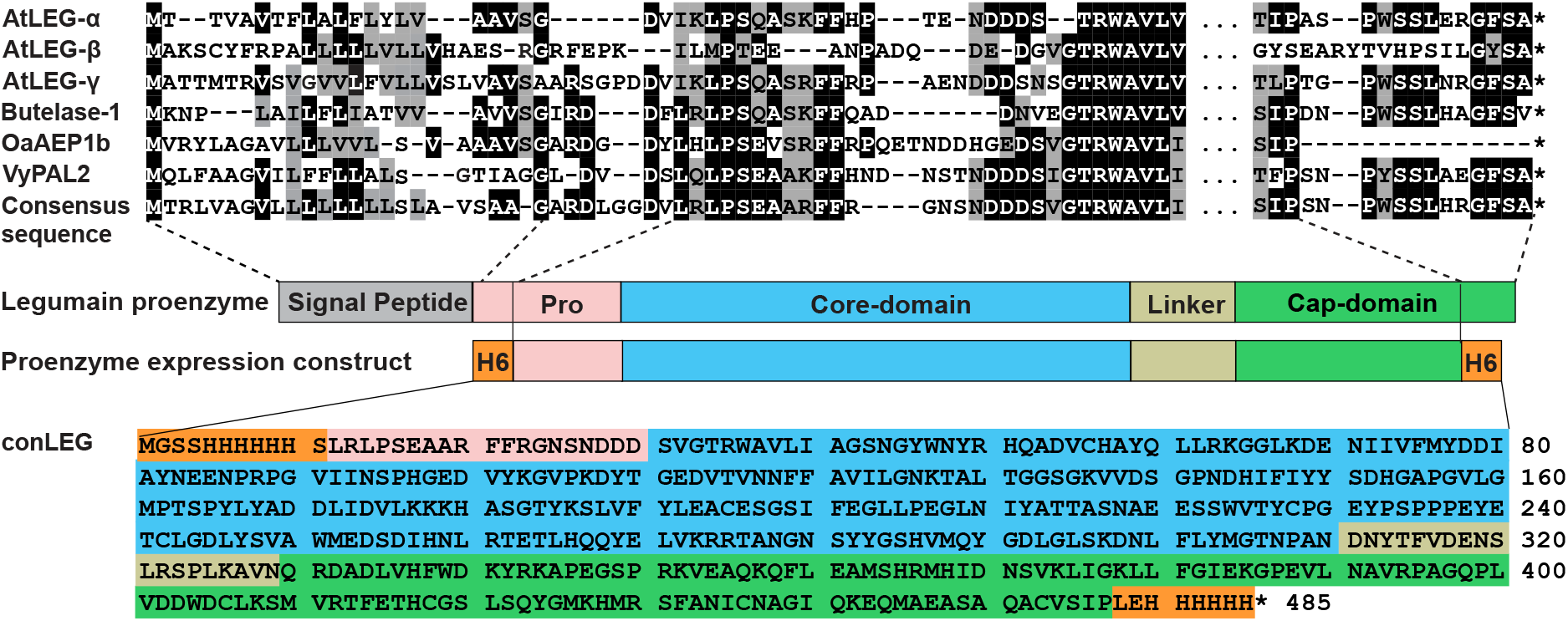
Construction of conLEG sequence for recombinant expression. The construct encoded full-length proenzyme derived from the consensus sequence of 1,500 plant legumains, with a His6-tag attached to the N-terminus of the conserved region of the pro-domain.

Recombinant protein was expressed from the conLEG construct as a proenzyme in *E. coli* Shuffle-T7B. After 48 h induction, 36-40 mg/L purified protein was obtained (Figure S2A). Acid-induced autoactivation of conLEG was carried out under different pHs and temperatures. The optimized conditions for autoactivation were acidification of a 1 mg/mL proenzyme solution to pH 4.0 and incubation for 2 h at 37 °C (Figure S2B). After size-exclusion chromatography (SEC) purification, a final yield of 12 mg active enzyme per 1 L culture was obtained.

### conLEG functions mainly as a protease, but has cyclase and splicing activity toward some substrates

To test the innate enzymatic ability of a plant legumain to catalyze hydrolysis, splicing and ligation, we carried out a functional study of activated conLEG toward four substrates. These four peptide substrates, including a pair of short peptides GN10-GL (GISYKPAYL<u>N</u>GL, MW 1295 Da) and GD10-GL (GISYKPAYL<u>D</u>GL, MW 1296 Da), the cyclotide-mimicking GN14-SLAN (GISTKSIPPISYR<u>N</u>SLAN, MW 1918 Da), and SFTI(D/N)-HV (GRCTKSIPPICFP<u>N</u>HV, MW 1768 Da) that mimics the precursor of the cyclic trypsin inhibitor SFTI-1, were chemically synthesized. Each was reacted with conLEG at eight reaction pHs ranging from 4.5 to 8.0.

At acidic pH, conLEG acts as an AEP and hydrolyzed all four peptide substrates as monitored by HPLC. At neutral and slightly basic pH, conLEG displayed low ligase activity (Figure 3A). For example, at pH 5, >60% of the 12-residue peptide GN10-GL was hydrolyzed to the 10-residue GN10 (GISYKPAYLN, MW, 1125 Da) with <3% cGN10 (cyclo-GISYKPAYLN, MW, 1107 Da), resulting in a very low cyclization/hydrolysis (C/H) ratio of 0.04 (Figure 3B). In contrast, at pH 7.5, 19% of GN10-GL was hydrolyzed and 26% was cyclized, yielding a C/H ratio of 1.4. Similar product distribution profiles in which the C/H ratio increased with pH were obtained for the other three substrates (Figure 3C-E).

**Figure 3.**
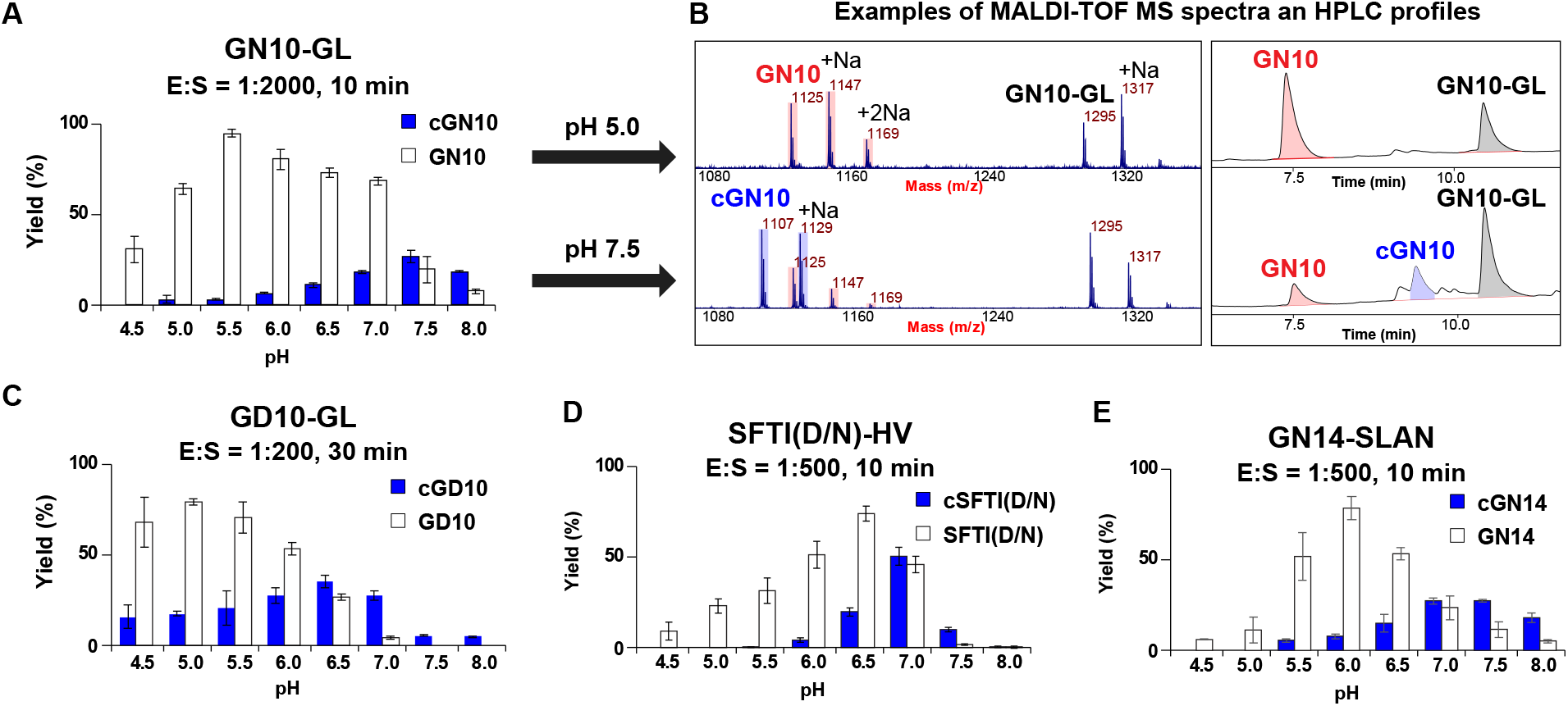
conLEG-mediated hydrolysis and cyclization of peptide substrates (A, B) GN10-GL, (C) GD10-GL, (D) SFTI(D/N)-HV and (E) GN14-SLAN. Reactions were carried out in triplicate and monitored by MALDI-TOF MS and quantified by HPLC. Representative examples of MS and HPLC data for the GN10-GL reaction at pH 5 and pH 7.5 are displayed in the panel on the right.

We also compared the catalytic effect of conLEG on two different peptide substrates: GN10-GL and GD10-GL, which have an Asn and Asp at the P1 recognition site and P1 position, respectively. P1-Asp resulted in higher C/H ratios at all eight pH values tested (Figure 3C). The generality of cyclic product formation in conLEG-mediated reactions suggest that plant legumains function intrinsically as AEPs under acidic pH conditions but the activity can also be bi-directional and influenced by extrinsic factors such as pH as well as by substrate and sequence.

To determine whether conLEG can act as a splicing enzyme when Asp instead of Asn is present at the P1 position, we synthesized a model substrate AI<u>N</u>GLRRGYSGS<u>D</u>ALEG (1735 Da) containing both Asn and Asp to mimic the precursor of the trypsin inhibitor McoTI-II (Heitz et al., 2001; Liew et al., 2021). At pH 5.0, we observed formation of the Asn-cleaved intermediate GLRRGYSGS<u>D</u>-ALEG (1437 Da) and a cyclic product cyclo(GLRRGYSGSD) (1049 Da), suggesting that conLEG could catalyze both Asn-specific hydrolysis and Asp-specific transpeptidation under acidic pH (Figure 4A). Under neutral and basic conditions, conLEG also catalyzed Asn-specific ligation (Figure 2A). A similar trimodal enzymatic activity was reported for McPAL1, the bioprocessing enzyme that mediates cyclization of McoTI-II at Asp via a splicing mechanism (Liew et al., 2021).

**Figure 4.**
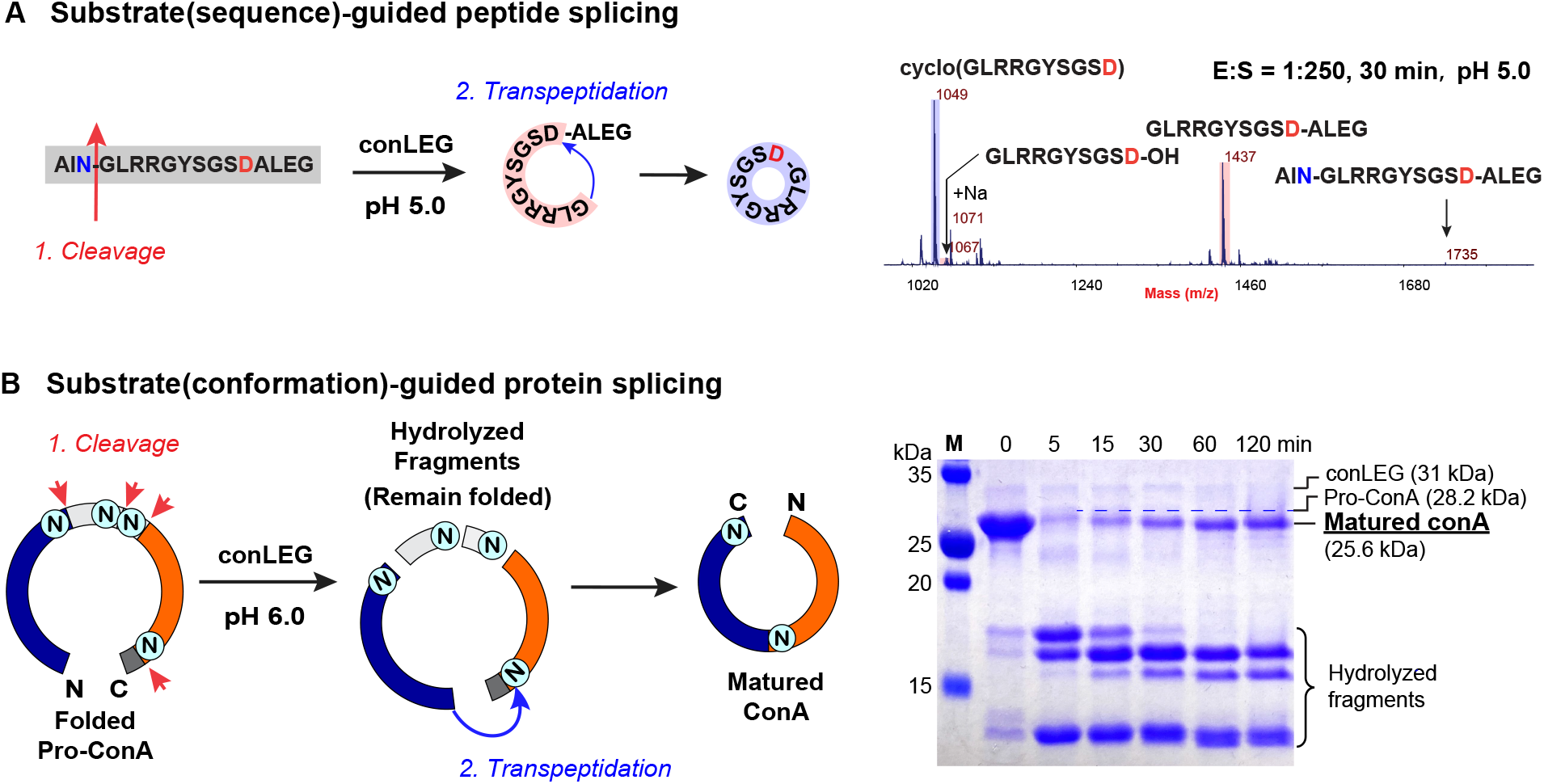
conLEG-mediated Asn/Asp-specific cleavage and transpeptidation. (A) conLEG-mediated N-terminal Asn-specific hydrolysis and C-terminal Asp-specific transpeptidation of AINGLRRGYSGSDSLEG to produce cyclo(GLRRGYSGSD). The reaction was carried out at pH 5.0 to facilitate release of the N-terminal tripeptide cap AIN. (B) conLEG-mediated maturation of conA by Asn-specific hydrolysis followed by structural-guided transpeptidation.

To show conLEG splicing activity toward proteins, we recombinantly expressed a protein substrate, concanavalin A precursor (pro-conA), which was recombinantly expressed, folded by dialysis, and then matured by conLEG-mediated-processing (Figure S3). Monitoring of conLEG splicing at different time points over 2 h using SDS-PAGE showed that folded pro-conA (28.2 kDa) was rapidly cleaved into smaller fragments within 5 min followed by gradual accumulation of mature conA (25.6 kDa) over time (Figure 4B). These results were similar to those reported for conA maturation with its native bioprocessing enzyme CeAEP1 (Nonis et al., 2021). Of note, the *in vitro* splicing reaction or transpeptidation reaction was slow and took hours to reach completion.

### Conversion of conLEG to conPAL1-3 by mutation of LAD residues in PAL substrate binding pockets

To convert conLEG to a PAL, we targeted ligase-activity determinants (LADs) in substrate binding pockets that determine whether the legumain activity is primarily AEP or PAL (Hemu et al., 2019; Hemu et al., 2020). In PALs such as butelase-1 and VyPAL-2, the critical direction-steering LAD motifs have hydrophobic residues substituted for two conserved Gly residues that flank the S1 catalytic center (Hemu et al., 2019; Nguyen et al., 2014). In conLEG, LAD1 and LAD2 have Gly225 and Gly155, respectively, which are both AEP-like. To convert conLEG into PAL-like ligases, we made a Gly225Val mutation in LAD1 to generate conPAL1 that mimics butelase-1, and a mutation Gly155Ala in LAD2 to generate conPAL2 that mimics VyPAL2 (Figure 5A). Finally, we engineered conPAL3, having a G155A/G225V double mutation in the substrate-binding pockets, which combines features of the two single mutants conPAL1 (butelase-1-like) and conPAL2 (VyPAL-like).

**Figure 5.**
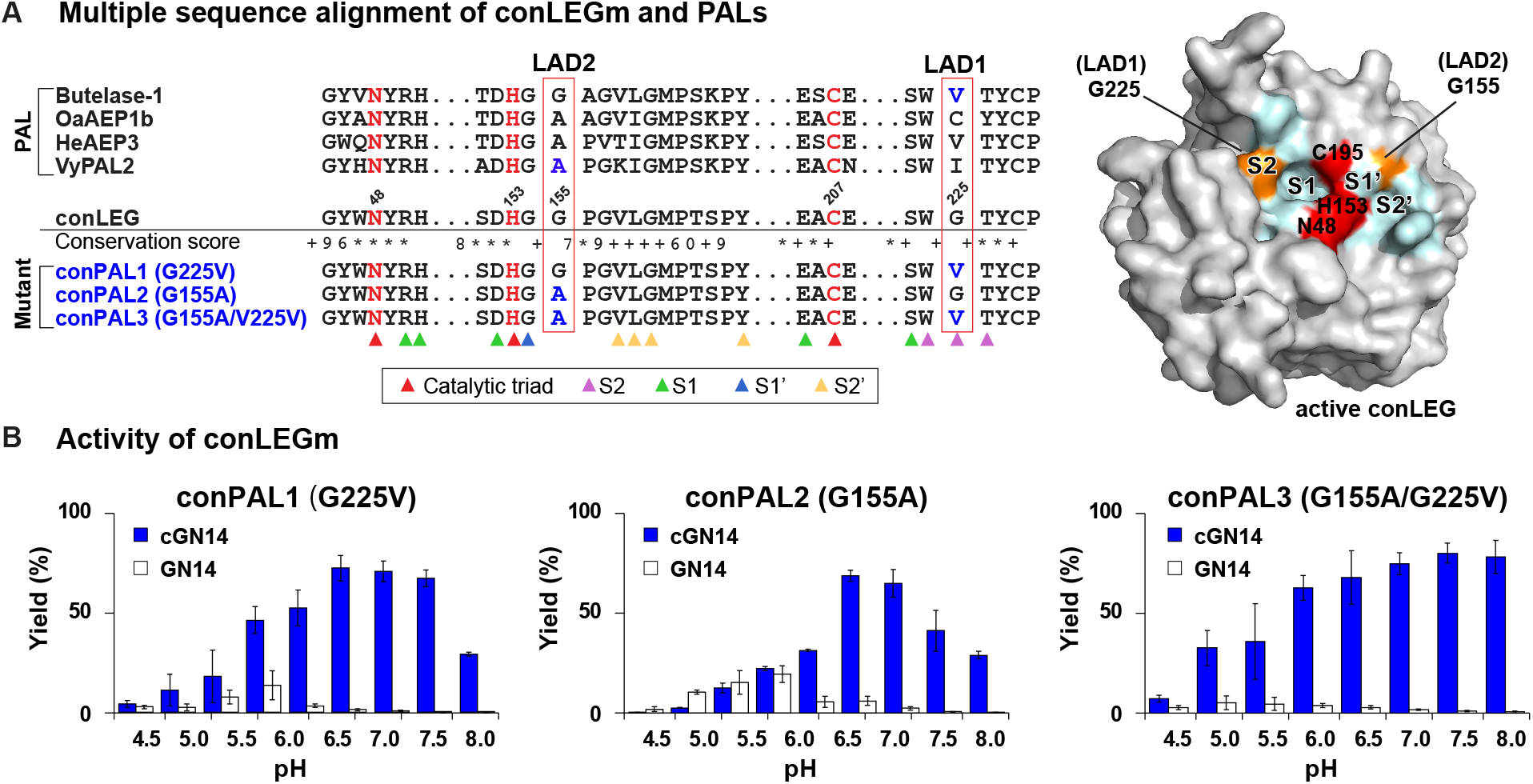
PAL-like conLEG mutants based on LAD hypothesis. (A) Location of the key LAD residues G225 and G155 in the conLEG and PAL sequence alignment and conLEG model structure. The catalytic triad N-H-C is in red. LAD-mutations in conPAL1-3 are highlighted in blue and orange in the model structure. Substrate binding residues S2-S1-S1’-S2’ are indicated by different arrows and colored cyan in the structure. (B) Activity screening of conPAL1-3 at different pHs with GN14-SLAN as the substrate.

Recombinant conPAL1-3 were expressed using the same protocol as for conLEG and their enzymatic activity was examined by MALDI-TOF MS at 25 °C with the substrate GN14-SLAN at a fixed molar ratio of E:S = 1:1000 and at eight different pHs, ranging from 4.5 to 8.0 (Figure 5B, Figure S4). In contrast to the AEP-like conLEG, all three conPALs displayed predominant ligase activity. In particular, the double mutant conPAL3 had a C/H ratio >20 (equivalent to >95% yield) at pH ≥6.5.

### conPAL3 displays high catalytic efficiency

We further characterized conPAL3 for its substrate specificity, catalytic efficiency, and stability. We first determined the optimal reaction conditions for conPAL3 by examining the initial catalytic rate of GN10-GL cyclization at three different pHs (6.0, 6.5 and 7.0) and seven different temperatures (10, 20, 25, 30, 37, 42 and 50 °C) (Figure 6). The conPAL3-mediated reactions had an optimal pH of 7.0 and the optimal reaction temperature was 25 °C. This result differs slightly from natural PALs like butelase-1 or VyPAL2, which have an optimal reaction temperature between 37 °C and 42 °C and an optimal pH of 6-6.5.

**Figure 6.**
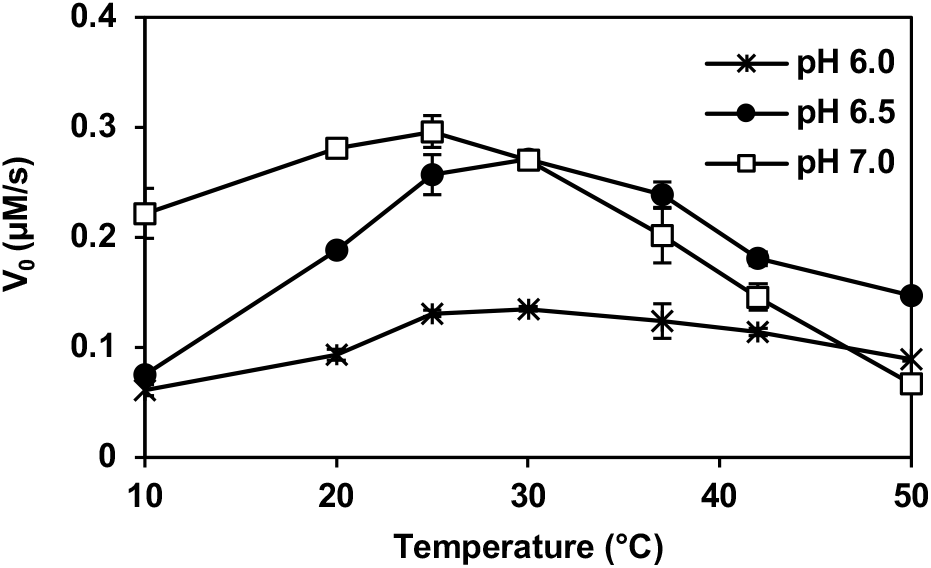
Optimal pH and temperature of conPAL3-mediated GN10-GL cyclization. Reactions were performed at seven different temperatures and three different pH values with a 1:10,000 E:S ratio. To calculate the initial catalytic rates, samples of the reaction mixtures taken every 30 s were quenched for quantitative analysis.

The amino acid preference of substrate-binding pockets S2-S1-S1’-S2’ was screened using panels of synthetic substrates having Asn or Asp at the P1 position and saturated variation at the P2, P1’, P2’, P1”, or P2” positions (Table 1A, Figure S5). The screening indicated that the conPAL3 S1 pocket favors Asn over Asp (Figure S5A). The catalytic efficiency against GN10-GL was at least 80-fold higher (*k*_cat_/K_m_ = 518,031 M^-1^s^-1^ at the optimum pH 7.0) relative to GD10-GL (*k*_cat_/K_m_ = 6,431 M^-1^s^-1^ at the optimum pH 5.0) (Figure S6A&B). The S2 pocket accepted any amino acid except the backbone-hindering Pro (Figure S5B). The S1’ pocket is the binding site for both the leaving group P1’ residue and the P1” residue from the incoming nucluophile. S1’ had broad amino acid tolerance to P1’ but was less tolerent to changes at the incoming P1”. Gly was the most favored P1” residue (Figure S5C). The S2’’ pocket, which also binds both P2’ and P2’’ residues, favored hydrophobic residues like Phe, Leu, Ile, and Met. This pocket is more tolerant to variations at P2’ than at incoming P2’’. Overall, conPAL3 has similar substrate preference to PALs but the less favorable leaving groups and incoming groups increased the likelihood of hydrolysis. Based on the results of the substrate-specificity screening, a model substrate GFSYKPAYSN-GI (MW 1303.44 Da) was designed and synthesized. At the optimum pH 7.0, catalytic efficiency of conPAL3 toward this model substrate was highly efficient at 2,289,260 M^-1^s^-1^, which is nearly 2-fold faster than the reported efficiency for butelase-1 (Table 1B, Figure S6C).

**Table 1.**
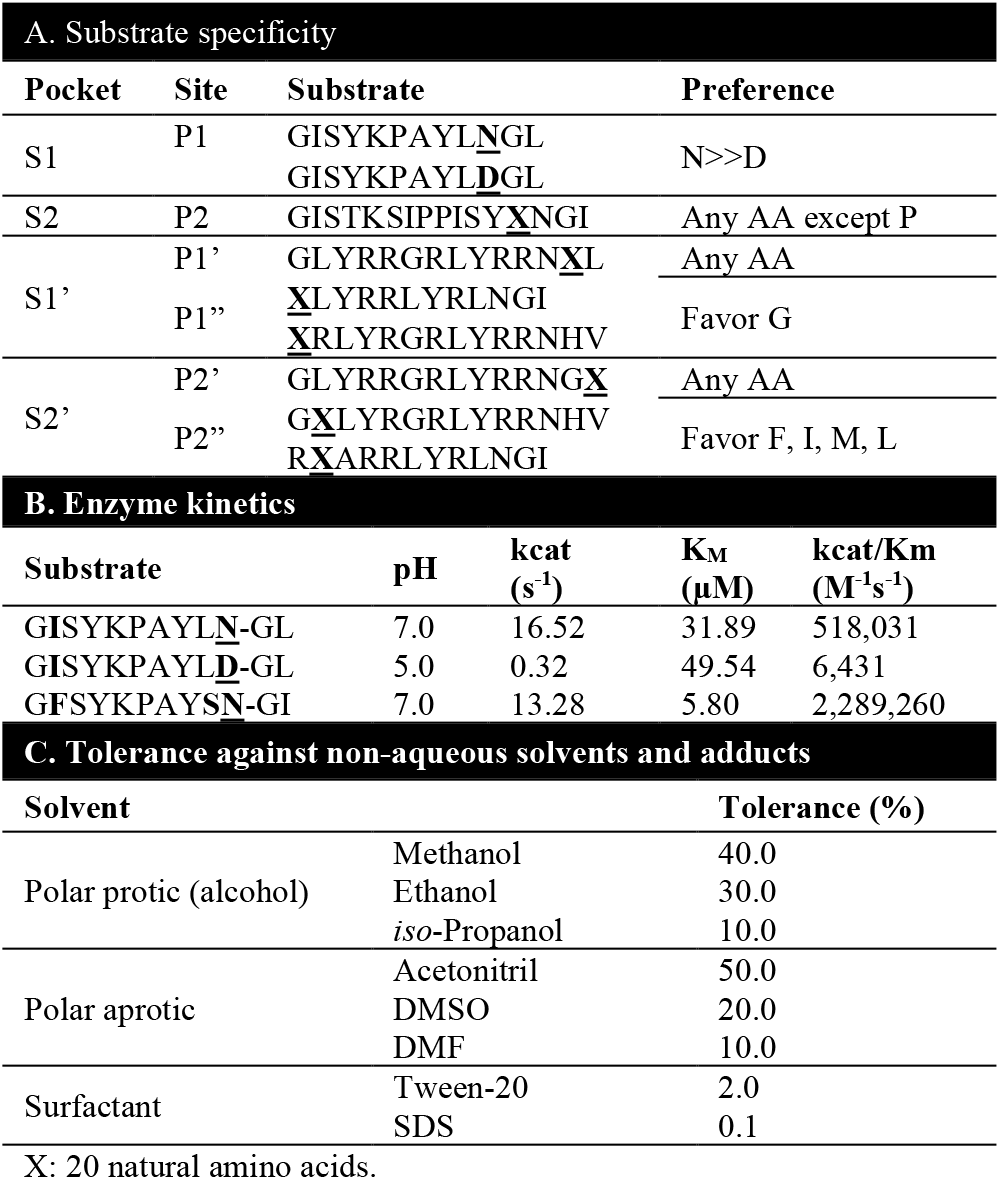
Enzymatic and chemical properties of conPAL3.

The tolerance of conPAL3 for non-aqueous solvents was examined by activity tests performed with various concentrations of polar protic solvents, polar aprotic solvents, and surfactants. conPAL3 retained 90% activity in the presence of 50% ACN, 40% MeOH, 30% EtOH10% iPrOH30% DMSO, 10% DMF, and 2% Tween-20, but was rapidly inactivated by 0.1% SDS (Table 1C, Figure S7).Substrate specificity

## Discussion

In this study, we combined a consensus AEP sequence found in 1,500 legumains with conserved ligase activity determinants (LADs) found in the substrate-binding sites of a small group of known PALs to generate new PALs having significantly improved biochemical properties (Figure 7). The direct use of a consensus sequence derived from a large set of plant legumain sequences reduced the phylogenetic bias. Compared with natural legumains expressed in the most economically-efficient bacteria system, we observed a 20-fold increase in conLEG expression, from an average of 2 mg/L proenzyme to about 40 mg/L. This increase may be due in part to improved mRNA stability and peptide solubility that facilitates translation and folding processes (Sternke et al., 2019). This high-yield and efficient conPAL3 could serve as a new model for PAL engineering and applications.

**Figure 7.**
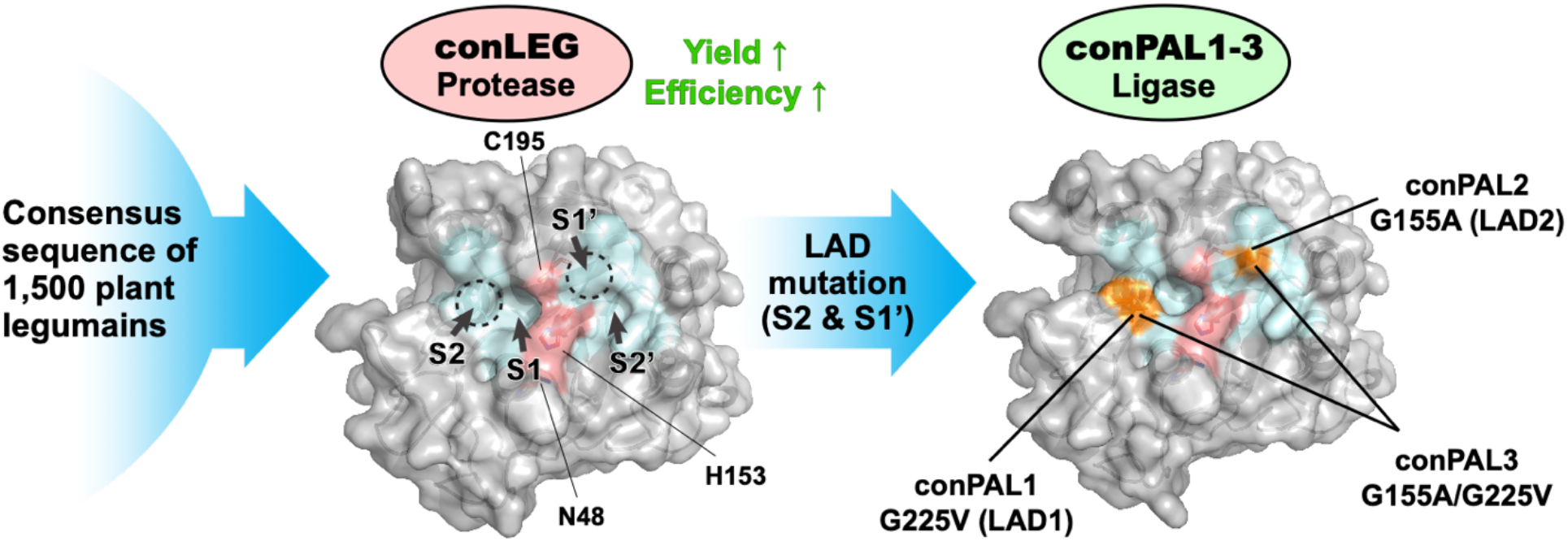
Schematic illustration of consensus design and LAD-based engineering of high-yield and efficient PALs.

The designed conLEG shares 64-88% core-domain identity with 20 previously described AEPs and PALs (Figure S8). The consensus design described here also improved the activity of plant legumains. The double mutant conPAL3 carrying LAD mutations on both sides (S2, and S1’-S2’) of the S1 Asx-binding pocket exhibited similar catalytic efficiency toward a model peptide as the fastest natural PAL, butelase-1. In addition, conPAL3 has broad substrate specificity toward P1-flanking sequences and optimal ligase activity at neutral pH and ambient temperature, as well as a wide tolerance to various non-aqueous solvents. We also observed that the reaction pH affects the thermal stability of conLEG, suggesting that the stability of its active conformation is closely associated with intramolecular charge-charge interactions. Together, our results demonstrate that a consensus design based on a large sequence dataset is a feasible approach to provide improved functional proteins.

Protease activity was the first identified and most frequently observed property of legumains. As such, the term AEP is often used interchangeably with legumain as a representative of the C13 subfamily. However, our work used an “averaged” sequence to show that plant legumain activity is intrinsically hydrolytic under acidic conditions, but can be conditionally bidirectional. The activity is influenced by environmental factors such as pH, substrate composition (such as P1-Asp) and conformation (such as folded conA), or by intrinsic factors such as LAD mutations that led to the emergence of PALs. Based on comparison of experimental results, we conclude that the selected intrinsic and extrinsic factors could exert profound influence on the enzymatic directionality toward AEPs or PALs.

## Materials and Methods

### Generation of plant legumain consensus sequence

Full amino acid sequences of butelase-1, OaAEP1b, AtLEGα-γ, and butelase-2 were used as representative plant PALs and AEPs to search homologous sequences in NCBI (BLASTp – non-redundant protein (nr) database, tBLASTx – nr and Transcriptome Shotgun Assembly (TSA) database) and 10KP database. The search yielded 1,500 non-duplicated hits as reported recently (Hemu et al., 2022). Multiple sequence alignment (MSA) was performed using the ClustalOmega program and analyzed by Jalview to generate visualization data including sequence Logo, consensus sequence, conservation, and occupancy. Regions having < 10% occupancy were removed from the consensus sequence.

### Plasmid construction and site-directed mutagenesis

The conLEG cDNA sequence was codon-optimized, synthesized and subcloned into the pET28a(+) vector with restriction cleavage by NdeI/XhoI to carry both N- and C-terminal His6 tags (Genscript). Primers carrying a G155A or G225V mutation were designed for site-directed mutagenesis using a Q5® Site-Directed Mutagenesis Kit (New England Biolabs).

### Recombinant expression of conLEG and mutants

Plasmids were transformed into SHuffle T7 *E. coli* competent cells (NEB, C3029J). LB-broth (kanamycin+) cultures (1 L) were grown at 30 °C with shaking at 180 rpm until the OD_600_ reached 0.4. The cultures were cooled to 16 °C before Isopropyl β-D-1-thiogalactopyranoside (IPTG, 0.1 mM) was added and the cultures were further incubated for 24─48 h. Cell pellets were harvested by centrifugation at 6,000 x g for 15 min at 4 °C and resuspended in cold lysis buffer (50 mM Na HEPES, 0.1 M NaCl, 1 mM EDTA, 5 mM β-mercaptoethanol, 0.1% TritonX-100, pH 7.5) in a ratio of 20 mL per 1 g pellet. Cells were lysed by sonication (10 min for 50 mL, 2s on/ 8s off) on ice. Clarified cell lysates were obtained by centrifugation at 12,000 x g for 15 min and subjected to affinity purification on a 5 mL Excel HisTrap affinity purification column (GE Life Sciences) equilibrated with IMAC-A buffer (20 mM sodium phosphate buffer pH 7.5, 0.1 M NaCl, 5% glycerol, 1 mM EDTA, 5 mM β-ME). The column was then washed with a mixture of IMAC-A and IMAC-B (0.5 M imidazole in buffer IMAC-A, pH 7.5) buffer containing 0.25 M imidazole (95/5 v/v, IMAC-A/IMAC-B). Targeted proteins were eluted with IMAC-B buffer containing 0.5 M imidazole. Fractions containing the targeted proenzymes were further purified on a size exclusion chromatography column (HiLoad 16/600 Superdex 200, Cytiva) equilibrated with SEC-7.5 buffer (20 mM sodium phosphate buffer pH 7.5, 0.1 M NaCl, 5% glycerol, 1 mM EDTA, 5 mM β-ME) to remove imidazole.

### Activation of conLEG and mutants

Acid-induced enzyme activation was performed at pH 3.5-6.0. Optimal activation was observed with pH 4.0-4.2 and incubation for 2-2.5 h at 37 °C with approximately 1 mg/mL proenzyme in 20 mM sodium citrate buffer containing 5 mM β-ME and 1 mM EDTA. Activated enzymes were purified by size exclusion chromatography equilibrated with SEC-4 buffer (20 mM sodium citrate buffer, 5 mM β-ME, 5% glycerol, 0.1 M NaCl, pH 4.0).Fractions containing target proteins were neutralized with storage buffer (20 mM sodium citrate, 5 mM β-ME,1 mM EDTA, 20% sucrose, 0.1 M NaCl, pH 5.5) and kept at 4 °C or stored at -80 °C after rapid freezing in liquid nitrogen.

### Preparation of peptide substrates

Peptide substrates (GN10-GL, GD10-GL, GN14-SLAN, SFTI(D/N)-HV) were synthesized by Fmoc chemistry using an automated microwave-synthesizer (Liberty Blue, CEM). Peptide substrate libraries (GISTKSIPPISY<u>X</u>NGI, <u>X</u>LYRRLYRLNGI, RXARRLYRLNGI, GLYRRGRLYRRN<u>X</u>L, <u>X</u>RLYRGRLYRRN-HV, GLYRRGRLYRRNG<u>X</u>, GLYRGRLYRRNHV, X = 20 natural amino acids) were purchased from GL Biochem.

### Preparation of recombinant pro-conA and conLEG-mediated maturation of conA

cDNA encoding His6-TEV-pro-conA sequence was synthesized and sub-cloned into pET28a(+) in framed with the N-terminal His-tag (Genscript), followed by transformation into Rosetta PLysS competent cells (Novagen). The overnight culture was amplified from 1 mL to 1 L with LB broth at 37 °C with shaking at 180 rpm until the OD_600_ reached 0.5. Recombinant expression of pro-conA was induced by addition of 0.5 mM IPTG and incubation for 24 h at 16 °C with shaking at 180 rpm. Cell pellet was harvested by centrifugation at 6,000 x g for 10 min at 4 °C. The pellets were resuspended in 10x w/v conA lysis buffer (20 mM MOPS, 0.5 M NaCl, 1 mM CaCl_2_, 1 mM MnCl_2_, 0.1% Triton X-100, pH 7). After 5 min sonication on ice, inclusion bodies were collected by centrifugation at 12,000 x g for 15 min at 4 °C. The pellets were washed twice with conA buffer (conA lysis buffer without Triton X-100). Purified inclusion bodies were resuspended in 40 mL refolding solution (conA buffer with 6 M guanidine-HCl, pH 7) followed by 1 min sonication. The soluble portion was isolated by centrifugation at 12,000 x g for 30 min at 4 °C. Recombinant protein was refolded by gradual dilution with 30x conA buffer followed by purification using size-exclusion chromatography. To remove the N-terminal tag, TEV protease was mixed with recombinant His6-TEV-pro-conA at a 1:10 w/w ratio and incubated at 25 °C for 4 h or 4 °C overnight. Pro-conA was purified from the TEV protease by reverse IMAC. For conLEG-mediated circular permutation, pro-conA was buffer-exchanged with legumain reaction buffer (20 mM sodium phosphate, 5 mM 2-mercaptoethanol, 1 mM EDTA, pH 6.0) using a centrifugal concentrator (Vivaspin Turbo 15, Sartorius) and concentrated to a final concentration of 1 mg/mL. Active conLEG was added to the conA solution to reach a molar ratio of conLEG : pro-conA = 1:100. Reactions were performed at 25 °C and monitored by SDS-PAGE at different time points.

### Functional studies of active conLEG and mutants

Functional studies of conLEG and mutants were carried out to examine intramolecular cyclization of peptide substrates at 37 °C in reaction buffers having a pH ranging from 4 to 8 (20 mM citrate or phosphate buffers with 1 mM EDTA and 5 mM β-mercaptoethanol). The peptide substrates GN12-GL (GLYRRGRLYRRN-GL) and SFTI1-GL (GRCTKSIPPICFPD-GL) were synthesized using Fmoc chemistry on an automated synthesizer (Liberty Blue, CEM) and purified by preparative RP-HPLC. Concentration of peptide substrates in peptide cyclization reactions was fixed at 20 μM unless specifically mentioned. Concentration of the activated conPAL3 used in the kinetic studies was 10 nM. All reactions were monitored by MALDI-TOF mass spectrometry (5800 Applied Biosystem, USA) and quantitatively analyzed by RP-HPLC on a C18 analytical column (Aeris™ WIDEPORE, Phenomenex, USA) after quenching with 1:1 v/v acetonitrile containing 0.1% trifluoroacetic acid.

## Author contributions

Xinya Hemu, Conceptualization, Methodology, Data curation, Formal analysis, Validation, Visualzation, Writing - original draft, review and editing; Xiaohong Zhang, Data curation, Formal analysis, Validation, Writing - review and editing; Hong Yi Chang, Data curation, Formal analysis, Validation, Writing - review and editing; Poh Jin En, Data curation, Formal analysis, Validation; James P. Tam Conceptualization, Resources, Supervision, Funding acquisition, Writing - original draft, review and editing.

