## Supplement Figures S1-S8 for "Consensus design and engineering of an efficient and high-yield Peptide Asparaginyl Ligase"

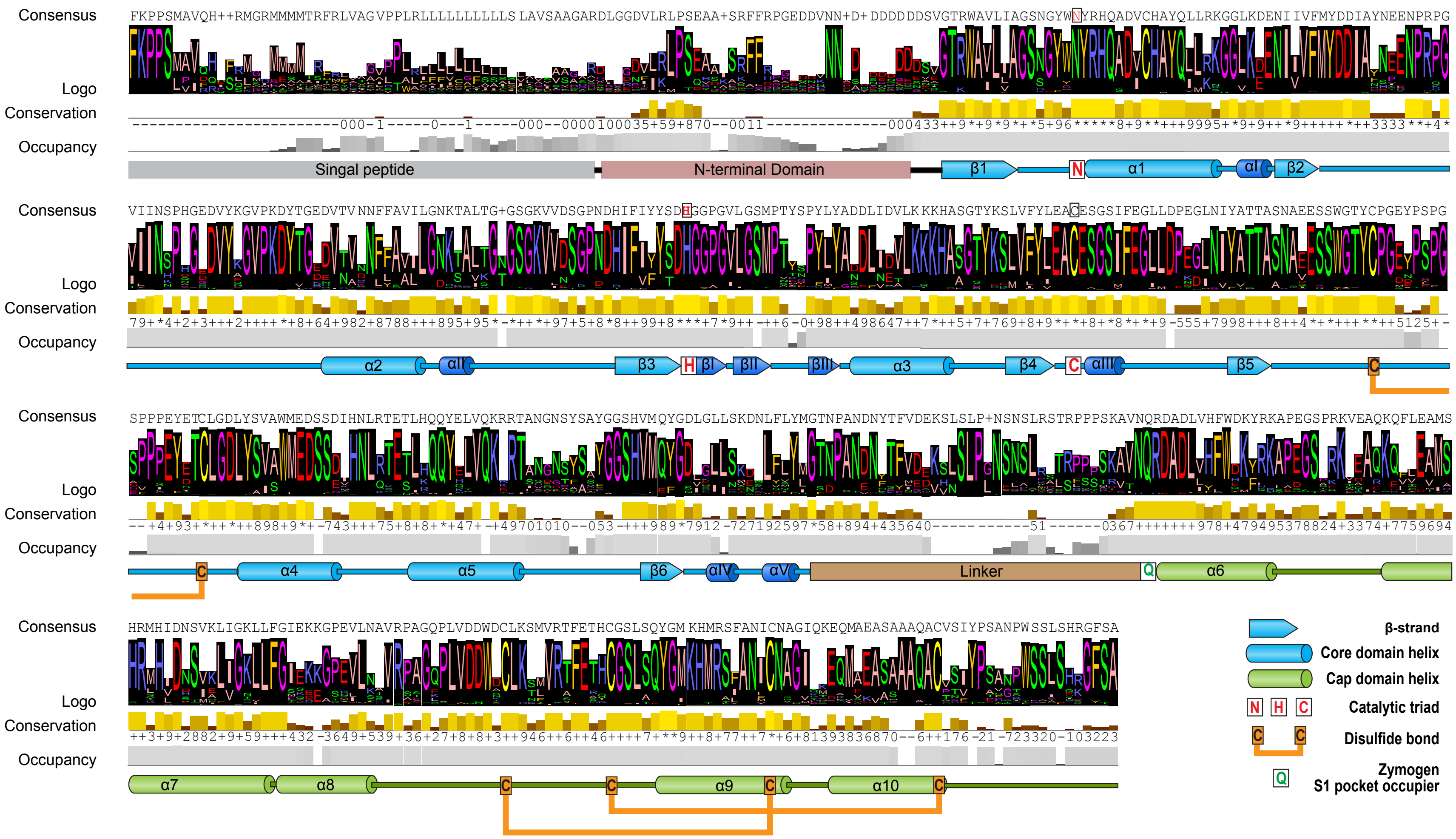


**Figure S1.** Construct and design of the consensus sequence of plant legumains for recombinant expression based on multiple sequence alignment and analysis of 1495 plant legumains. The secondary structures, protein domains and functional features were annotated based on crystal structure of AtLEGβ (PDB code: 5NIJ).

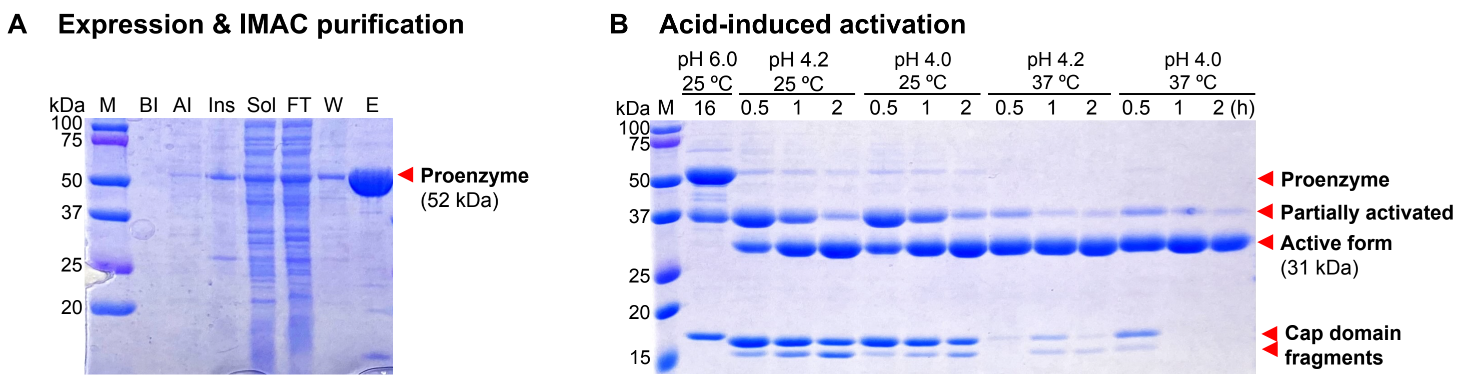


**Figure S2**. Recombinant expression of conLEG. (A) SDS-PAGE of extraction and immobilized metal affinity chromatography (IMAC) purification. M, protein marker. BI, before induction. AI, after induction. Ins, insoluble lysate. Sol, soluble extract. FT, IMAC flow-through. W, IMAC wash-through. E, IMAC elution. (B) SDS-PAGE of activation conditions.

TEV protease cleavage site

His6-TEV-Pro-conA MGHHHHHHSSGGTENLYFQ-S$_{119}$THETNALHFMFNQFSKDQKDLILQGDATTGTDGNLELTRVSSNGSPQGSSVGRALFYAPV
HIWESSAVVASFEATFTFLIKSPDSHPADGIAFFISNIDSSIPSGSTGRLLGLFPDAN$_{237}$VIRNSTTIDFNAAYNA$_{1}$DTIVAVE
LDTYPNTDIGDPSYPHIGIDIKSVRSKKTAKWNMQNGKVGTAHIIYNSVDKRLSAVVSYPNADSATVSYDVDLDNVLPEWVRVG
LSASTGLYKETNTILSWSFTSKLKSN$_{118}$EIPDIATVV-

Pro-conA S$_{119}$THETNALHFMFNQFSKDQKDLILQGDATTGTDGNLELTRVSSNGSPQGSSVGRALFYAPVHIWESSAVVASFEATFTFLIK
SPDSHPADGIAFFISNIDSSIPSGSTGRLLGLFPDAN$_{237}$VIRNSTTIDFNAAYNA$_{1}$DTIVAVELDTYPNTDIGDPSYPHIGIDI

conLEG cleavage sites

KSVRSKKTAKWNMQNGKVGTAHIIYNSVDKRLSAVVSYPNADSATVSYDVDLDNVLPEWVRVGLSASTGLYKETNTILSWSFTS
KLKSN$_{118}$EIPDIATVV-

conLEG ligation site

Matured conA A$_{1}$DTIVAVELDTYPNTDIGDPSYPHIGIDIKSVRSKKTAKWNM
QNGKVGTAHIIYNSVDKRLSAVVSYPNADSATVSYDVDLDNVL
PEWVRVGLSASTGLYKETNTILSWSFTSKLKSN$_{118}$S$_{119}$THETN
ALHFMFNQFSKDQKDLILQGDATTGTDGNLELTRVSSNGSPQG
SSVGRALFYAPVHIWESSAVVASFEATFTFLIKSPDSHPADGI
AFFISNIDSSIPSGSTGRLLGLFPDAN$_{237}$

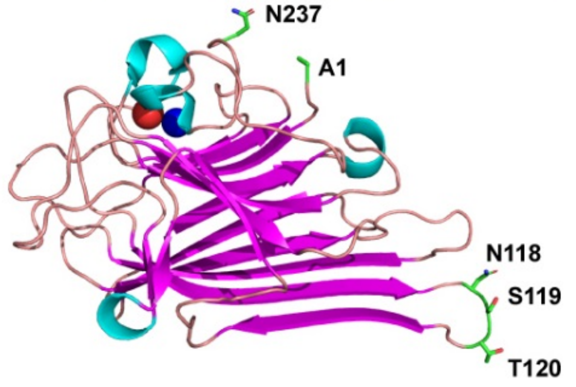


**Figure S3.** Amino acid sequences of His6-TEV-Pro-conA, pro-conA precursor and matured conA. The legumain-processing sites, A1, N118, S119, N237 were labelled on the structure of a matured conA monomer (PDB ID: 1NLS).

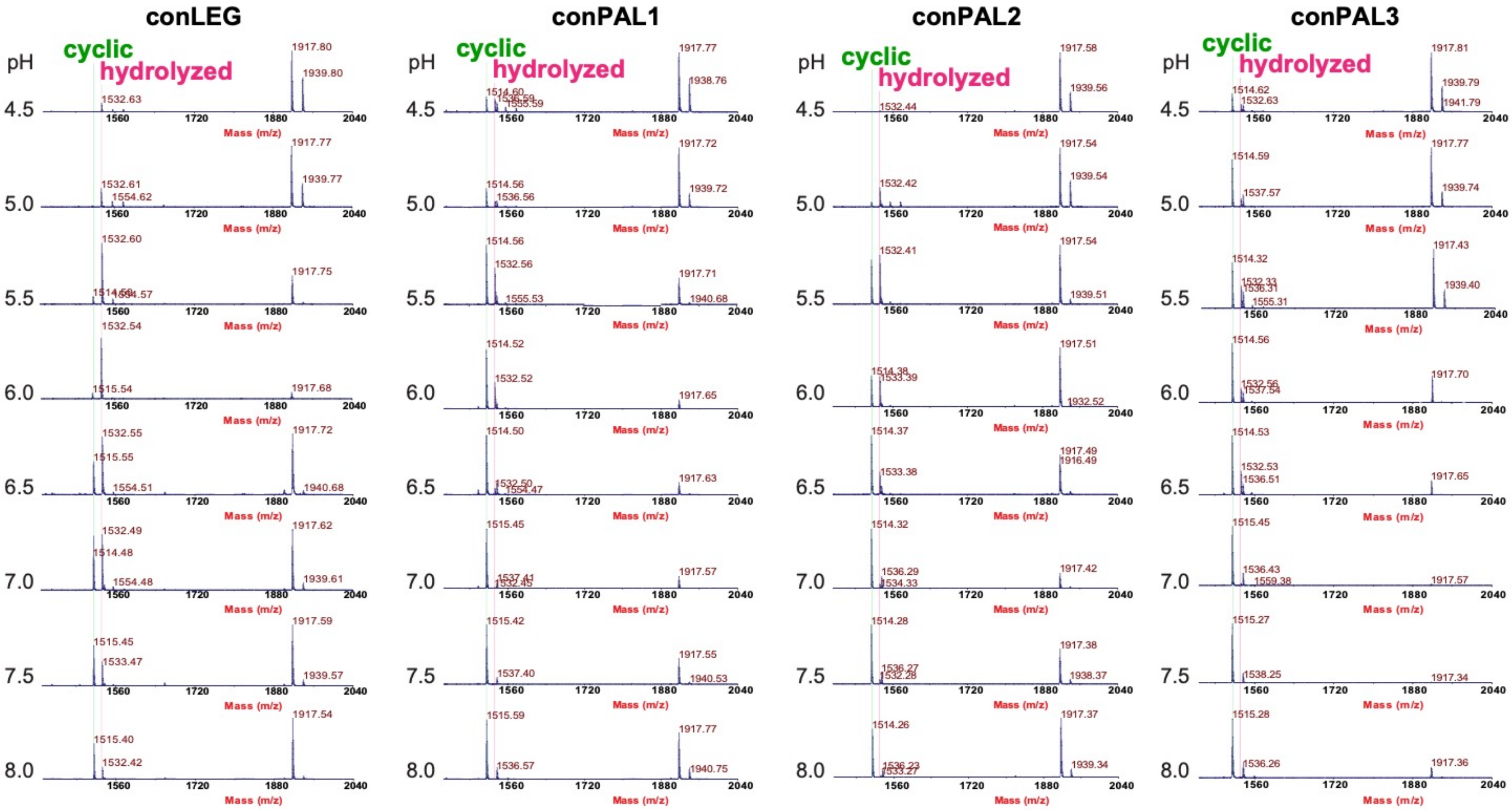


**Figure S4**. Examples of MS data for conLEG1-3 activity screening.

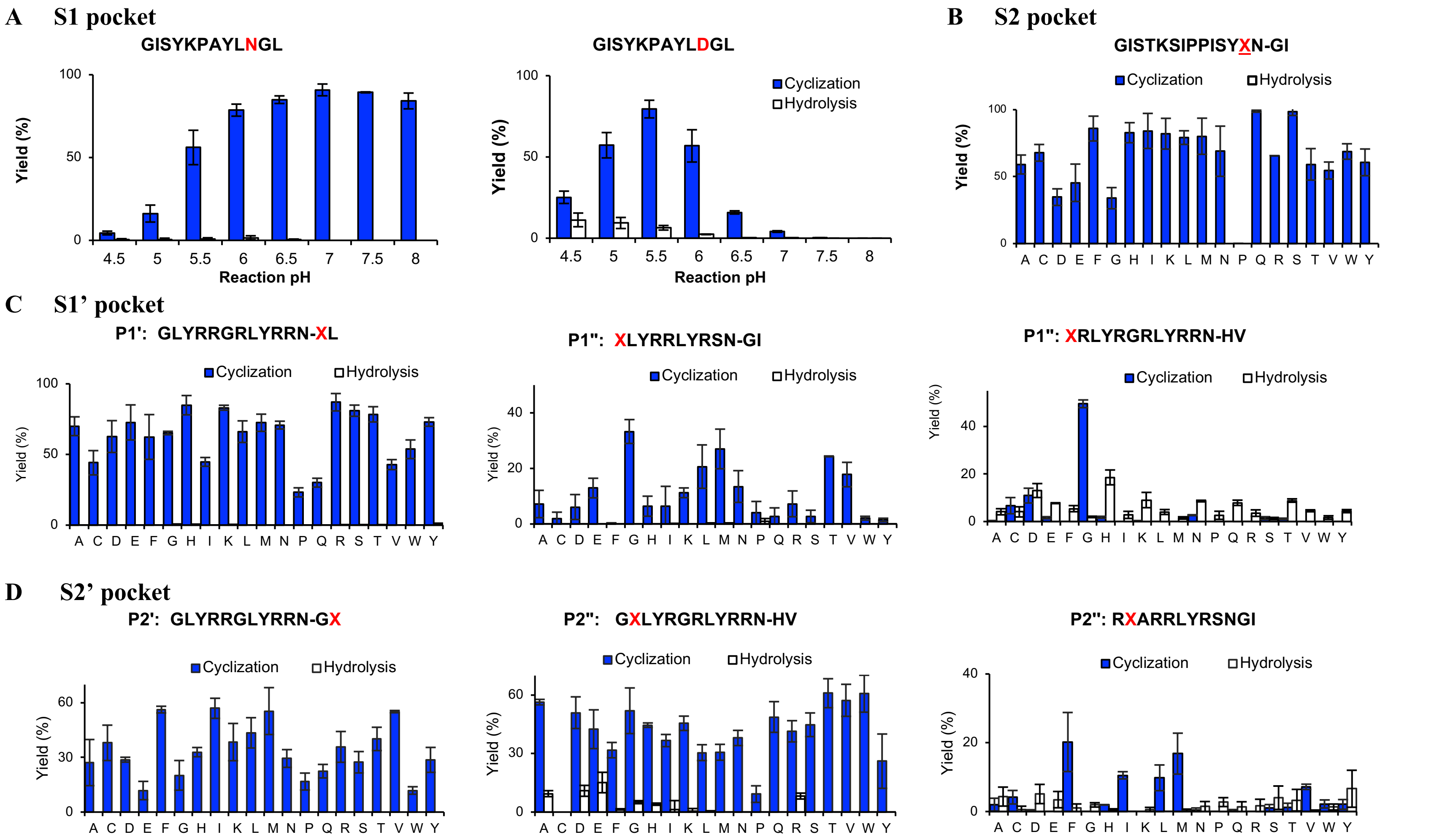


**Figure S5.** Binding specificity of S2-S1-S1'-S2' substrate pockets of conPAL3. (A) Specificity of S1 pocket against Asn and Asp. (B) Specificity study of S2 pocket shows that S2 pocket has broad tolerance to any residues except Pro. (C) Specificity study of S1' pocket against P1' and P1'' residues shows that S1' pocket is not highly selective but favor P1''-Gly in ligation reactions. (D) Specificity study of S2' pocket against P2' and P2'' residues shows that S2' pocket favors hydrophobic residues including Phe, Ile, Leu, Met, and Val. But in the case that P1'/P1'' is occupied by Gly, P2' or P2'' couould be any residues.

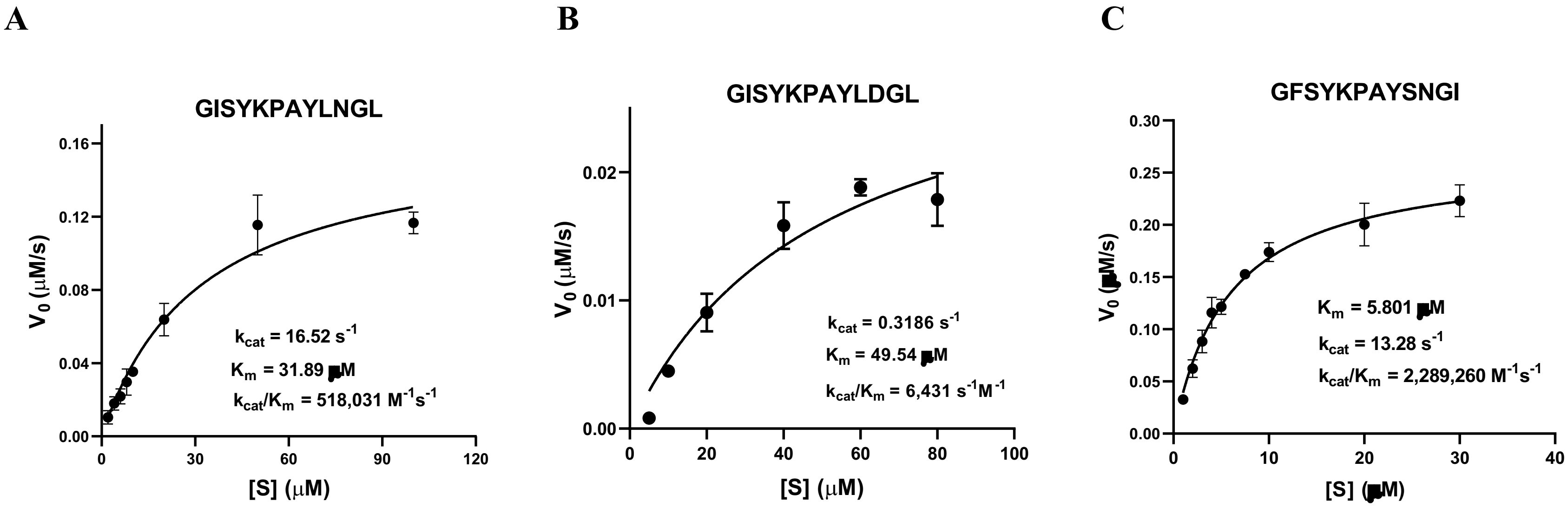


**Figure S6.** Enzyme kinetic study of conPAL3 in three peptide cyclization reactions. Reactions were performed at pH 5.0 for GISYKPAYLDGL and pH 7.0 for the other two substrates at 25 °C in triplicates. Reaction products were quantified by UPLC.

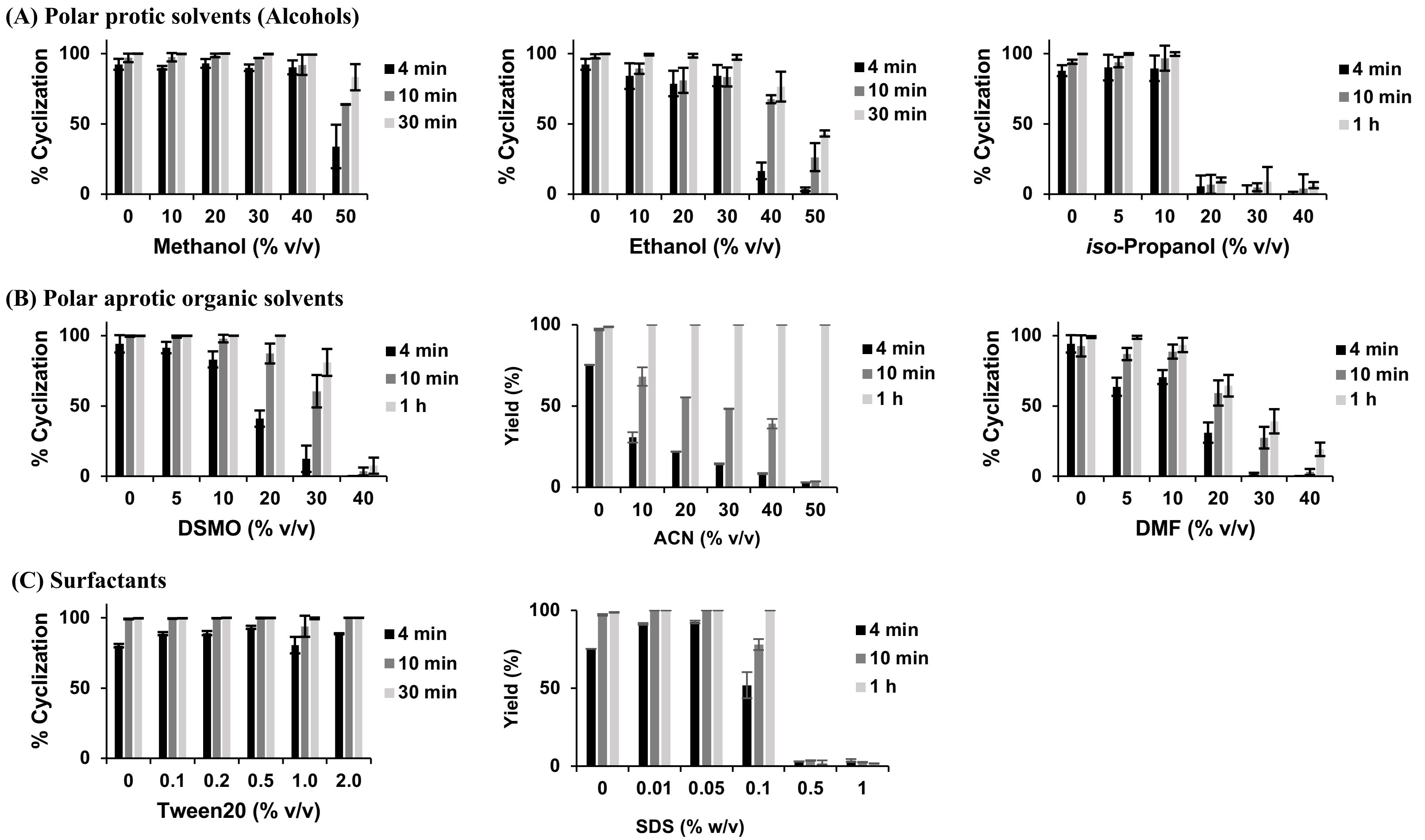


**Figure S7. Tolerance against non-aqueous solvents.** (A) Polar protic solvents: methanol, ethanol, isopropanol. (B) Polar aprotic organic solvents: DMSO, ACN, DMF. (C) Surfactants: (non-ionic) Tween-20 and (anionic) SDS.

```
PAL
Butelase1     GTRWAVLVAGSKGYVNYRHQADVCHAYQILKKGGLKDENIIVFMYDDIAYNESNPHPGVIINHPYGSDVYKGVPKDYVGEDINPPNFYAVLLANKSALTGTGSGKVLDSGPNDHVFIYYTDHGGAGVLGMPSKPYIAASDLNDVLKKKHASGTYKSIVFYVESCESGSMFDGLLPEDHNI   180
OaAEP1b       GTRWAVLIAGSKGYANYRHQAGVCHAYQILKRGGLKDENIVVFMYDDIAYNESNPRPGVIINSPHGSDVYAGVPKDYTGEEVNAKNFLAAILGNKSAITG-GSGKVVDSGPNDHIFIYYTDHGAAGVIGMPSKPYLYADELNDALKKKHASGTYKSLVFYLEACESGSMFEGILPEDLNI   179
OaAEP3        GTRWAVLIAGSKGYDNYRHQADVCHAYQILKRGGLKDENIVVFMYDDIAYNESNPRPGVIINSPHGSDVYAGVPKDYTGDEVNAKNFLAAILGNKSAITG-GSGKVVDSGPNDHIFIYYTDHGAPGVIGMPSKPYLYADELNDALRKKHASGTYKSMVFYLEACEAGSMFDGLLPDGLNI   179
OaAEP4        GTRWAVLLAGSNGYWNYRHQADLCHAYQILKRGGLKDENIVVFMYDDIAYNEENPRPGVIINSPHGSDVYAGVPKDYTGDEVNAKNFLAAILGNKSAITG-GSGKVVDSGPNDHIFIYYTDHGAPGVIGMPSKPYLYADELNDALRKKHASGTYKSMVFYLEACEAGSMFDGLLPDGLNI   179
OaAEP5        GTRWAVLIAGSKGYDNYRHQADVCHAYQILKRGGLKDENIVVFMYDDIAYNESNPRPGVIINSPHGSDVYAGVPKDYTGEEVNAKNFLAAILGNKSAITG-GSGKVVDSGPNDHIFIYYTDHGAAGVIGMPSKPYLYADELNDALKKKHASGTYKSLVFYLEACEAGSMFEGLLTDDLNI   179
HeAEP3        GTRWAVLIAGSKGWQNYRHQADVCHAYQILKKGGLKDENIIVFMYDDIAYNESNPRPGIVINKPKGEDVYKGVPKDYTGENVNAVNFLAVLLANRSALTG-GSGKVLDSGPNDRIFIYYTDHGAPVTIGMPSKPYLVAKDLVDTLKKKHAAGTYKSMVFYIESCESGSMFDGLLPEDANI   179
VyPAL1        GTRWAVLIAGSKGYHNYRHQADVCHMYQILRKGGVKDENIIVFMYDDIAYNESNPFPGIIINKPGGENVYKGVPKDYTGEDINNVNFLAAILGNKSAIIG-GSGKVLDTSPNDHIFIYYADHGAPGKIGMPSKPYLYADDLVDTLKQKAAAGTYKSMVFYVEACNAGSMFEGLLPEGMNI   179
VyPAL2        GTRWAVLIAGSKGYHNYRHQADVCHMYQILRKGGVKDENIIVFMYDDIAYNESNPFPGIIINKPGGENVYKGVPKDYTGEDINNVNFLAAILGNKSAIIG-GSGKVLDTSPNDHIFIYYADHGAPGKIGMPSKPYLYADDLVDTLKQKAATGTYKSMVFYVEACNAGSMFEGLLPEGTNI   179
VyPAL4        GTTWAVLIAGSKGYHNYRHQADVCHMYQILRKGGVKDENIIVFMYDDIAYNESNPFPGIIINKPGGENVYKGVPKDYTGEDINNVNFLAAILGNKSAIIG-GSGKVLDTSPNDHIFIYYADHGAPGKIGMPSKPYLYADDLVDTLKQKAAAGTYKSMVFYVEACNAGSMFEGLLPEGMNI   179
VyPAL5        GTTWAVLIAGSKGYHNYRHQADVCHMYQILRKGGVKDENIIVFMYDDIAYNESNPFPGIIINKPGGENVYKGVPKDYTGEDINNVNFLAAILGNKSAIIG-GSGKVLDTSPNDHIFIYYADHGAPGKIGMPSKPYLYADDLVDTLKQKAATGTYKSMVFYVEACNAGSMFEGLLPEGTNI   179

AEP
Butelase2     GTRWAVLVAGSNGYENYRHQADVCHAYQLLIKGGLKEENIVVFMYDDIAWHELNPRPGVIINNPRGEDVYAGVPKDYTGEDVTAENLFAVILGDRSKVKG-GSGKVINSKPEDRIFIFYSDHGGPGVLGMPNEQILYAMDFIDVLKKKHASGGYREMVIYVEACESGSLFEGIMPKDLNV   179
VyAEP1        GTRWAVLVAGSNGFGNYRHQADVCHAYQLLRKGGLKEENIVVFMYDDIATNQLNPRPGIIINHPQGEDVYHGVPKDYTGAEVNAHNLYAVLLGDKSAVKG-GSGKVVNSKPDDRIFVYYSDHGGPGVLGMPNLPYVYAMDFIDTLKKKHAANSYREMVVYVEACESGSLFEGVMPKDLNI   179
OaAEP2        GTRWAVLLAGSNGYWNYRHQADLCHAYQILKRGGLKDENIVVFMYDDIAYNEENPRPGVIINSPHGSDVYAGVPKDYTGDQVNAKNFLAAILGNKSAITG-GSGKVVNSGPNDHIFIYYTDHGGPGVLGMPVGPYIYADDLIDTLKKKHASGTYKSLVFYLEACESGSMFEGLLPEGLNI   179
CeAEP         GTRWAVLVAGSNGYGNYRHQADVCHAYQLLIKGGVKEENIVVFMYDDIAYNAMNPRPGVIINHPQGPDVYAGVPKDYTGEDVTPENLYAVILGDKSKVKG-GSGKVINSNPEDRIFIFYSDHGGPGVLGMPNAPFVYAMDFIDVLKKKHASGGYKEMVIYIEACESGSIFEGIMPKDLNI   179
RcAEP         GTRWAVLVAGSMGFGNYRHQADVCHAYQLLRKGGLKEENIIVFMYDDIAKNELNPRPGVIINHPQGEDVYAGVPKDYTGEHVTAKNLYAVLLGDKSAVQG-GSGKVVDSKPNDRIFLYYSDHGGPGVLGMPNLPYLYAMDFIEVLKKKHAAGGYKKMVIYVEACESGSIFEGIMPKDVDI   179
HaAEP         GTKWAVLVAGSKGYGNYRHQADVCHAYQVLKKGGLKDENIVVFMYDDIAKSEMNPRPGIIINSPKGEDVYAGVPKDYTGKNVTVDNLSAVLLGDRSAVKG-GSGKVVDSKPEDRIFLFYSDHGGPGVLGMPNEPHLVAKDLVDVLKKKHAMGTYKEMVIYLEACESGSIFEGILPEDLNI   179
AtVPE-alpha   STRWAVLVAGSSGYWNYRHQADVCHAYQLLKKGGVKEENIVVFMYDDIAKNEENPRPGVIINSPNGEDVYNGVPKDYTGDDVNVDNLLAVILGNKTAVKG-GSGKVVDSGPNDHIFIYYSDHGGPGVLGMPTSPYLYANDLNDVLKKKHASGTYKSLVFYLEACESGSIFEGLLPEGLNI   179
AtVPE-beta    GTRWAVLVAGSSGYGNYRHQADVCHAYQILRKGGLKEENIVVLMYDDIANHPLNPRPGTLINHPDGDDVYAGVPKDYTGSSVTAANFYAVLLGDQKAVKG-GSGKVIASKPNDHIFVYYADHGGPGVLGMPNTPHIYAADFIETLKKKHASGTYKEMVIYVEACESGSIFEGIMPKDLNI   179
AtVPE-gamma   GTRWAVLVAGSSGYWNYRHQADICHAYQLLRKGGLKEENIVVFMYDDIANNYENPRPGTIINSPHGKDVYQGVPKDYTGDDVNVDNLFAVILGDKTAVKG-GSGKVVDSGPNDHIFIFYSDHGGPGVLGMPTSPYLYANDLNDVLKKKHALGTYKSLVFYLEACESGSIFEGLLPEGLNI   179
AtVPE-delta   GTRWAVLVAGSNEYYNYRHQADICHAYQILRKGGLKDENIIVFMYDDIAFSSENPRPGVIINKPDGEDVYKGVPKDYTKEAVNVQNFYNVLLGNESGVTG-GNGKVVKSGPNDNIFIYYADHGAPGLIAMPTGDEVMAKDFNEVLEKMHKRKKYNKMVIYVEACESGSMFEGILKKNLNI   179

conLEG        GTRWAVLIAGSNGYWNYRHQADVCHAYQLLRKGGLKDENIIVFMYDDIAYNEENPRPGVIINSPHGEDVYKGVPKDYTGEDVTVNNFFAVILGNKTALTG-GSGKVVDSGPNDHIFIYYSDHGGPGVLGMPTSPYLYADDLIDVLKKKHASGTYKSLVFYLEACESGSIFEGLLPEGLNI   179
              .* ****:***  : ******.:** **:* :**:*:***:*:******    ** ** :** * * :** ******.   :.  *:  .:*.:.. : * *.***: : *:*.:*::*:***.   :.**    : * :: :.*.:      *..:*.*:*:*::**:*:*:: .. ::

                                                                                                                                   %Identity(conLEG)
PAL
Butelase1     YVMGASDTGESSWVTYCPLQHPSPPP--EYDVCVGDLFSVAWLEDCDVHNLQTETFQQQYEVVKNKTIVAL-IEDGTHVVQYGDVGLSKQTLFVYMGTDPANDN   281 | 73.48
OaAEP1b       YALTSTNTTESSWCYYCPAQEN-PPP-PEYNVCLGDLFSVAWLEDSDVQNSWYETLNQQYHHVDKRIS------HASHATQYGNLKLGEEGLFVYMGSNPANDN   275 | 74.55
OaAEP3        YALTASNTTEGSWCYYCPGQ-DAGPP-PEYSVCLGDFFSIAWLEDSDVHNLRSETLNQQYHNVKNRISY------ASHATQYGDLKRGVEGLFLYLGSNPENDN   275 | 75.55
OaAEP4        YALTASNTTEGSWCYYCPGQ-DAGPP-PEYSVCLGDFFSIAWLEDSDVHNLRSETLNQQYHNVKNRISY------ASHATQYGDLKRGVEGLFLYLGSNPENDN   275 | 75.91
OaAEP5        YALTASNATEGSCPYYCPGDLNYSPP-PEYDVCLGDFFSIAWLEDSDVHNLRSETLNQQYHNVKNRISY------ASHATQYGDLKRGVEGLFLYLGSNPENDN   276 | 76.00
HeAEP3        YGMTATNSTEGSWVTYCPGQTDDYPEDDEYDVCFGDLWSVAWLEDCDAHNLRTETLDQQYEVVKKKIEY------AHIPAQYGNVSLAKDSLFVYMGTDPANDN   277 | 71.27
VyPAL1        YAMTASNSTEGSLIAYCAGVTPGVPL--EIVTCLGDLWSITFLEDCDAHNLRTETVHQQFELVKKKIAY------ASTVSQYGDIPISKDSLSVYMGTDPANDN   275 | 71.90
VyPAL2        YAMAASNSTEGSWITYCPGT-PDFPP--EFDVCLGDLWSITFLEDCDAHNLRTETVHQQFELVKKKIAY------ASTVSQYGDIPISKDSLSVYMGTDPANDN   274 | 72.89
VyPAL4        YAMTASNSTEGSLIAYCAGVTPGVPL--EIVTCLGDLWSITFLEDCDAHNLRTETVHQQFELVKKKIAY------ASTVSQYGDIPISKDSLSVYMGTDPANDN   275 | 71.53
VyPAL5        YAMAASNSTEGSWITYCPGT-PDFPP--EFDVCLGDLWSITFLEDCDAHNLRTETVHQQFELVKKKIAY------ASTVSQYGDIPISKDSLSVYMGTDPANDN   274 | 72.53

AEP
Butelase2     FVTTASNAQENSWGTYCPGTEPSPPP--EYTTCLGDLYSVAWMEDSESHNLRRETVNQQYRSVKERTSNFKDYAMGSHVMQYGDTNITAEKLYLFQGFDPATVN   281 | 71.33
VyAEP1        YVTTASNAQENSWGTYCPGEG--APP--EYNTCLGDLYSVAWMEDSESHNLKKEAIKDQYKTVKARTSDSSTYHSGSHVMEYGNRSIRAEKLYLYQGFDPATVN   279 | 68.71
OaAEP2        YATTASNAEESSWGTYCPGEYP-SPP-PEYDTCLGDLYSVAWMEDSEVHNLRSETLKQQYHLVKARTSNGN-SAYGSHVMQYGDLKLSVDNLFLYMGTNPANDN   280 | 87.50
CeAEP         YVTTASNAQENSFGTYCPGMNPPPPE--EYVTCLGDLYSVSWMEDSETHNLKRETVQQQYQSVRKRTSNSNSYRFGSHVMQYGDTNITAEKLYLYHGFDPATVN   281 | 71.33
RcAEP         YVTTASNAQESSWGTYCPGMEPSPPP--EFTTCLGDLYSVAWMEDSESHNLKKETVKQQYSSVKARTSNYNTYAAGSHVMQYGNQSIKADKLYLFQGFDPASVN   281 | 73.12
HaAEP         YATTASGAQENSYGTYCPGTEPSPPP--EYITCLGDLYSVAWMEDSETHNLKKESLEQQFNKVKKRTSNSNTYNTGSHVMEYGSKDIKPEKVYLYLGFDPATVN   281 | 71.33
AtVPE-alpha   YATTASNAVESSWGTYCPGEDP-SPP-SEYETCLGDLYSVAWMEDSDIHNLQTETLHQQYELVKKRTAGSG-KSFGSHVMEFGDIGLSKEKLVLYMGTNPANEN   280 | 86.79
AtVPE-beta    YVTTASNAQESSYGTYCPGMNPSPPS--EYITCLGDLYSVAWMEDSETHNLKKETIKQQYHTVKMRTSNYNTYSGGSHVMEYGNNSIKSEKLYLYQGFDPATVN   281 | 69.18
AtVPE-gamma   YATTASNAEESSWGTYCPGEEP-SPP-PEYETCLGDLYSVAWMEDSGMHNLQTETLHQQYELVKRRTAPVG-YSYGSHVMQYGDVGISKDNLDLYMGTNPANDN   280 | 88.21
AtVPE-delta   YAVTAANSKESSWGVYCPESYPPPPS--EIGTCLGDTFSISWLEDSDLHDMSKETLEQQYHVVKRRVGSD--VPETSHVCRFGTEKMLKDYLSSYIGRNPENDN   279 | 64.03

conLEG        YATTASNAEESSWGTYCPGEYP-SPP-PEYETCLGDLYSVAWMEDSDIHNLRTETLHQQYELVKRRTANGN-SYYGSHVMQYGDLGLSKDNLFLYMGTNPANDN   280
              :   ::.: *.*   **       *   *  .*.** :*::::**.  ::   *:..:*:  *  :             .:*     : :  : * :* . *
```

**Figure S8.** Sequence alignment and percentage of sequence identity between conLEG and 20 other plant legumains.
